# A novel hypothalamic-midbrain circuit for model-based learning

**DOI:** 10.1101/2023.03.02.530856

**Authors:** Ivy B. Hoang, Joseph J. Munier, Anna Verghese, Zara Greer, Samuel J. Millard, Lauren E. DiFazio, Courtney Sercander, Alicia Izquierdo, Melissa J. Sharpe

## Abstract

Behavior is often dichotomized into model-free and model-based systems ^1, 2^. Model-free behavior prioritizes associations that have high value, regardless of the specific consequence or circumstance. In contrast, model-based behavior involves considering all possible outcomes to produce behavior that best fits the current circumstance. We typically exhibit a mixture of these behaviors so we can trade-off efficiency and flexibility. However, substance use disorder shifts behavior more strongly towards model-free systems, which produces a difficulty abstaining from drug-seeking due to an inability to withhold making the model-free high-value response ^3–10^. The lateral hypothalamus (LH) is implicated in substance use disorder ^11–17^ and we have demonstrated that this region is critical to Pavlovian cue-reward learning ^18, 19^. However, it is unknown whether learning occurring in LH is model-free or model-based, where the necessary teaching signal comes from to facilitate learning in LH, and whether this is relevant for learning deficits that drive substance use disorder. Here, we reveal that learning occurring in the LH is model-based. Further, we confirm the existence of an understudied projection extending from dopamine neurons in the ventral tegmental area (VTA) to the LH and demonstrate that this input underlies model-based learning in LH. Finally, we examine the impact of methamphetamine self-administration on LH-dependent model-based processes. These experiments reveal that a history of methamphetamine administration enhances the model-based control that Pavlovian cues have over decision-making, which was accompanied by a bidirectional strengthening of the LH to VTA circuit. Together, this work reveals a novel bidirectional circuit that underlies model-based learning and is relevant to the behavioral and cognitive changes that arise with substance use disorders. This circuit represents a new addition to models of addiction, which focus on instrumental components of drug addiction and increases in model-free habits after drug exposure ^3–10^.

## Introduction

Decision-making is often governed by two types of cognitive strategies: model-free learning and model-based learning ^1, 2^. Model-free systems promote behavior by assigning reward-predictive states and responses a scalar value, making rewarded actions more likely to be repeated in the future. On the other hand, model-based agents develop chains of associations between states, responses, and rewarding outcomes, which facilitates evaluation of future outcomes and enables flexible decisions. A model-free approach can be helpful for fast, efficient decision-making. However, a model-based strategy is more advantageous when we need to carefully consider surrounding circumstances. This is particularly important when our wants and desires have negative consequences in the longer term. Imagine you’ve recently developed a gluten allergy. It’s important that you use this new information to avoid things like pastries, despite the fact that these items have a high value from your allergy-free past. The adaptive response to avoid pastries after discovering your allergy is facilitated by model-based decision-making, which allows you to contemplate the consequences of your choice beyond immediate gratification. Understanding the mechanisms involved in these different decision-making strategies is important because alterations in their balance has been consistently implicated in psychological disorders, particularly substance use disorder ^3–10^. For example, individuals with substance use disorder are thought to continue drug taking because they fail to contemplate the negative consequences of drug use in the long run, which is argued to reflect a shift towards model-free decision-making after drug exposure ^3–10^.

One brain region that contributes to reward-directed behavior and is implicated in substance use disorder is the lateral hypothalamus (LH) ^11–21^. Historically, the function of the LH has been limited to the innate motivation to consume rewards, especially in the context of feeding ^22–24^. Thus, a role for LH in substance use has been assumed to reflect an increased innate motivation for rewards, driving sustained drug-seeking ^13^. However, recently we have demonstrated that GABAergic neurons in LH (LH_GABA_) play a more complex role in behavior than previously thought. Specifically, we have revealed that LH is necessary to learn and express information about cues paired with rewards ^18, 19^, over and above a role in the innate drive to consume rewards. Importantly, the way in which LH contributes to reward learning seems to be in a manner that prioritizes learning of information most relevant to rewards ^18^. That is, while activity in LH_GABA_ neurons is necessary for learning about reward-paired cues, these neurons oppose learning about information not immediately relevant to rewards. This demonstrates that changes in LH activity could influence a balance in learning about reward-related and other information, suggesting that hypothalamic circuits constitute a dynamic learning system that can arbitrate between different types of learning in different scenarios. In this way, the strengthening of LH circuits reported after drug exposure ^11–13, 15–17^ could increase learning and behavior directed towards drug and other reward related cues, characteristic of substance use disorder and predictive of future drug-seeking ^25–28^.

While we know the LH is involved in reward learning and these circuits are strengthened in addiction ^11–13, 15–17^, there are many questions that remain unanswered. For example, the nature of reward learning acquired by the LH is unknown. This could involve detailed representations between cues and sensory-specific rewards, indicative of model-based learning. Alternatively, LH could help assign value to reward-paired cues, reflective of model-free learning. Further, the wider circuity supporting reward learning in the LH is unclear. One candidate for this is input from dopamine neurons in the ventral tegmental area (VTA), which we now understand to be capable of supporting both model-free and model-based associative learning ^29–47^. However, there are very few studies that have suggested the presence of a projection from dopamine neurons to VTA ^48, 49^, and none that explore the function of this projection. These questions are important in helping us understand how strengthening of LH circuits might contribute to substance use disorder. In particular, the manner in which Pavlovian drug cues influence addictive behavior is understudied relative to the vast amount of work that has been done characterizing how instrumental actions are altered by drug exposure. To this end, we were also interested in revealing how drug self-administration impacts learning and behavior directed to reward-paired cues, and whether this relates to changes in our hypothalamic-midbrain circuit.

## Results

### LH_GABA_ neurons are necessary for behavior governed by model-based associations between cues and rewards

We have previously shown that LH_GABA_ neurons are necessary for both the acquisition and expression of cue-reward associations ^18, 19^. However, it is unclear whether these cue-reward associations reflect the general, scalar value of the reward (i.e., model-free) or the unique, sensory-specific properties of the reward (i.e., model-based). To answer this question, we probed the content of cue-reward information in LH_GABA_ neurons (**Figure 1A**). We first infused GAD-Cre rats bilaterally with a Cre-dependent adeno-associated virus (AAV) carrying either the inhibitory halorhodopsin (AAV5-Ef1a-DIO-eNpHR3.0-eYFP; NpHR; *n*=9) or a control vector (AAV5-Ef1a-DIO-eYFP; eYFP; *n*=14) into LH and implanted optic fibers bilaterally above LH (**Figure 1B-C**). This would allow us to optogenetically inhibit LH_GABA_ neurons ^18, 19^. Four weeks after surgery, rats were food restricted and began Pavlovian conditioning procedures. Here, two distinct, 10-sec auditory cues (click and white noise; 8 sessions; 12 presentations/session) were presented with one stimulus leading to delivery of two 45-mg sucrose pellets (Test Diet, MA; CS+) and the other without consequence (CS-; counterbalanced). Across conditioning, eYFP and NpHR groups increased time spent in the food port during the CS+ relative to CS-presentation with no between-group differences (**Figure 1D**; CS+ vs. CS-: *F*_(1,21)_ = 63.483, *p*<0.001; session: *F*_(3,63)_ = 1.056, *p*=0.374; group: *F*_(1,21)_ = 0.728, *p*=0.403; CS x group: *F*_(1,21)_ = 0.891, *p*=0.356; session x group: *F*_(3,63)_ = 0.106, *p*=0.956; CS x session: *F*_(3,63)_ = 5.813, *p*=0.001; CS x session x group: *F*_(3,63)_ = 0.109, *p*=0.954).

**Figure 1.**
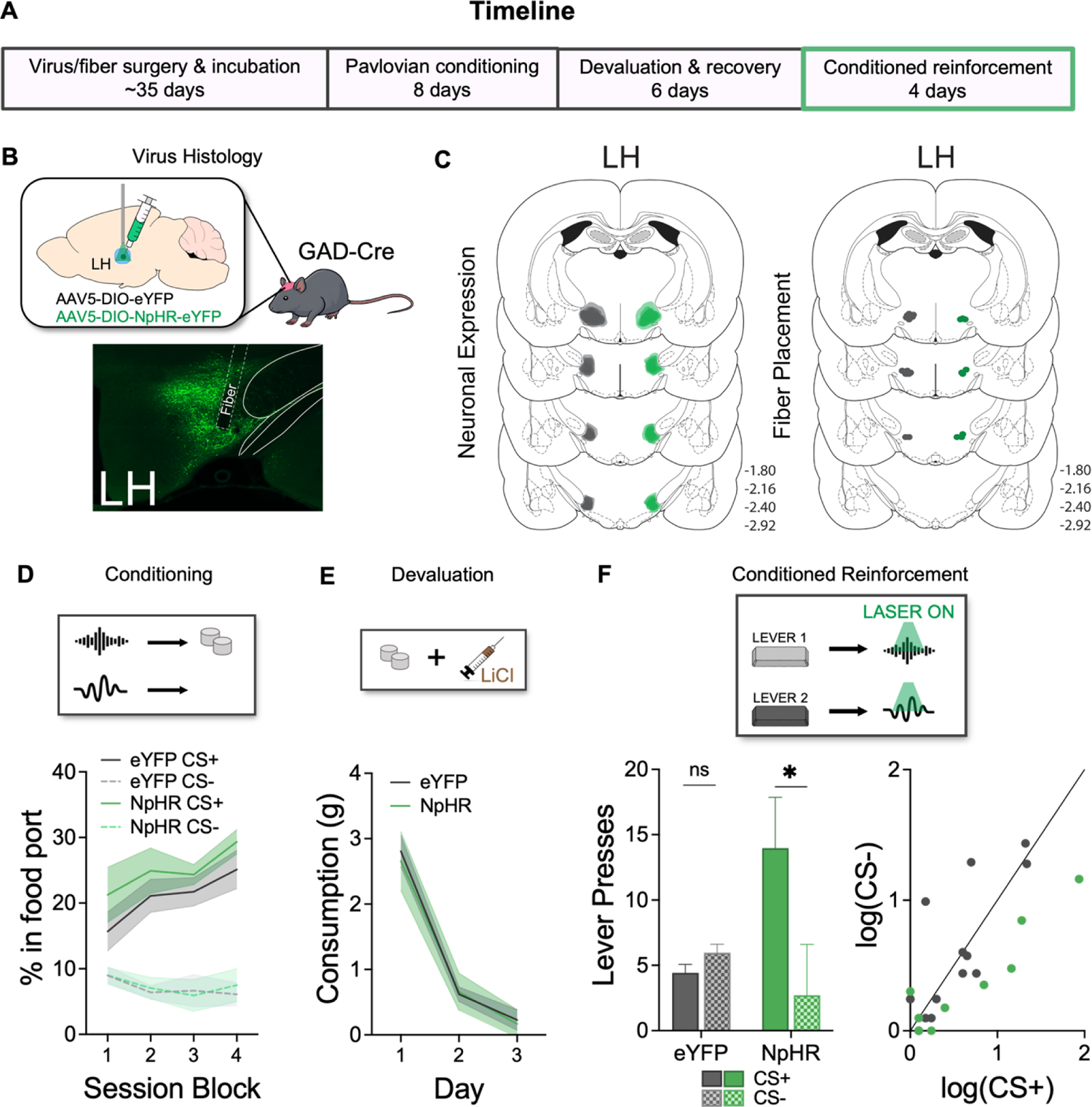
Inhibition of LH_GABA_ neurons prevents the use of model-based associations to guide behavior. (A) Experimental timeline. (B) Optogenetic approach: GAD-Cre rats were bilaterally infused with a Cre-dependent AAV with inhibitory halorhodopsin (NpHR; *n*=9), or a control virus (eYFP; n=14), in LH and implanted with optic fibers in LH. Below shows a unilateral example of bilateral virus expression in the cell bodies of GABAergic neurons in LH. (C) *Left*: Unilateral representation of bilateral virus expression in LH for the eYFP group (grey) and the NpHR group (green). *Right*: Dots indicate approximate location of fiber tips in LH. (D) Rats learned that one auditory cue led to food pellets (CS+), and another was without consequence (CS-). Responding is represented as the percent of time spent in the food port (mean ± SEM). Rats in both eYFP and NpHR groups increased responding to the CS+ relative to the CS-. (E) Rats received pairings of LiCl with the reward and all rats reduced consumption of the food reward. (F) Next, rats were allowed to press two levers to earn presentations of either CS+ or CS. Here, we inhibited LH_GABA_ neurons during presentation of CS+ and CS-. Rats in the eYFP group do not demonstrate conditioned reinforcement for the CS+ predicting the devalued reward. However, rats in the NpHR group showed robust conditioned reinforcement for the CS+. Rates of responding are represented as number of lever presses made for presentation of either cue (mean ± SEM). Individual data points reflect responding of each rat as a normalized value (logarithmic transformation) for eYFP (grey) and NpHR (green) groups. To the extent that responding for the CS+ and CS-is equal, dots should congregate on the diagonal. Rats in the eYFP group showed low levels of lever-pressing for both CS+ and CS-. However, NpHR rats showed high responding for the CS+ and not CS-. \**p* ≤ 0.05, mean (± SEM).

After learning about the CS+ and CS-, rats underwent a devaluation procedure, where the reward associated with CS+ was paired with injection of lithium chloride (LiCl; 0.15M, 10 mL/kg; 3 days). This procedure produced a taste aversion to the reward, reflected in a reduction in consumption of the reward across injection days (**Figure 1E**; day: *F*_(2,42)_ = 40.758, *p*<0.001; group: *F*_(1,21)_ = 0.038, *p*=0.846; day x group: *F*_(2,42)_ = 0.048, *p*=0.953). The devaluation procedure allows us to test the associative information contained in the cue-reward associations in LH. That is, if LH harbors model-based information, this association will be sensitive to reward devaluation. This is because the cue will evoke a representation of the reward, and the reward will evoke a feeling a sickness, which will lead the rat to reduce responding to the CS ^50–52^. However, if the information harbored in LH is model-free and based on a general value that has transferred to the CS across learning, the CS will not evoke a representation of the reward and responding will be insensitive to devaluation ^50, 53–55^. As inhibition of LH_GABA_ neurons during a cue will simply reduce the appetitive response ^18, 19^, we need to circumvent this issue by arranging a situation where we assess the content of information in LH_GABA_ without requiring a response during inhibition of LH_GABA_ neurons. Thus, instead of presenting the CS and measuring appetitive responding, we gave rats a test where they could press one lever to get presentation of the CS+ and another for the CS-. During this test, we delivered green light (532nm, 16-18mW) into the brain to inhibit LH_GABA_ neurons during CS+ and CS-presentation. This allowed us to selectively inhibit LH_GABA_ neurons during the CS *after* the response was made. As rats had no prior experience with the levers, continued lever-pressing would indicate that the CSs were capable of supporting development of instrumental response and were valuable in some way (known as the well-established phenomenon, conditioned reinforcement ^56–60^). However, given we had devalued the reward paired with the CS+, if rats are using model-based associative information to direct behavior, then this CS should not be capable of supporting conditioned reinforcement because the CS would be associated with the now devalued reward ^57, 61^. Indeed, we found that the eYFP group did not demonstrate the conditioned reinforcement effect after devaluation (**Figure 1F**). That is, there was no difference in the ability of the CS+ or CS- to drive conditioned reinforcement, reflecting sensitivity of this effect to devaluation ^57, 61^. In contrast, inhibition of LH_GABA_ neurons prevented the ability of rats in the NpHR group to use model-based information to drive behavior, illustrated by the ability of the CS+ to support conditioned reinforcement despite devaluation of the associated reward (**Figure 1F**). This was supported by statistical analyses which revealed no main effect of CS or group (CS+ vs. CS-: *F*_(1,21)_ = 2.615, *p*=0.121; group: F_(1,21)_ = 0.001, *p*=0.970), but a significant CS by group interaction (CS x group: *F*_(1,21)_ = 7.061, *p*=0.015). Simple-effect analyses following the interaction revealed no difference in lever-press responding for the CS+ and CS-in the eYFP group (*F*_(1,21)_ = 0.072, *p*=0.415). However, there was a significant difference in lever-press responding for the CS+ and CS- in the NpHR group (*F*_(1,21)_ = 0.294, *p*=0.012). These results demonstrate that LH_GABA_ neurons encode model-based associations that entail representations of the cue and the sensory-specific features of the predicted reward and so inactivating these neurons prevented the ability to use model-based associative information to govern responding.

### Confirming the presence of a novel dopaminergic projection from VTA to LH

Our first experiment provides evidence to suggest that the LH contains model-based information that can be called upon to influence adaptive behavior. This begs the question of which neural substrates facilitate this function. One candidate mechanism is input from dopamine neurons in the VTA ^35^. Historically, dopamine prediction errors have been thought to contribute to cue-reward learning by endowing cues with a model-free, scalar value ^29–34^. However, recent studies have revealed that these phasic signals can also support the development of model-based associations ^36–47^. For example, VTA_DA_ neurons are necessary for the development of associations between cues and sensory-specific representations of rewards ^38, 40^. While there is a body of literature showing dopamine activity regulates LH function, this is largely based on local pharmacological manipulations through dopamine-receptor agonists and antagonists and focused on the canonical role of LH in regulating feeding ^48, 62–66^. Thus, while the VTA is well-situated anatomically and functionally to contribute to model-based encoded in LH, the role of this circuit in learning is unknown.

Given there is sparse anatomical evidence for the existence of VTA projections to LH and few showing they are dopaminergic in nature ^48, 49, 67, 68^, we first verified the existence of VTA dopamine (VTA_DA_) input to LH by injecting retrograde tracer cholera toxin subunit B (CTb-555; Alexa Fluor^TM^ 555 conjugate) into LH (**Figure 2A**). Any neuronal projections terminating in LH take up the retrograde tracer and subsequently express in the originating cell bodies (**Figure 2C**). We then used an antibody to stain tyrosine hydroxylase (TH), an enzyme that converts tyrosine into dopamine, and imaged the VTA (**Figure 2D**). We found that injection of the CTb in LH resulted in considerable double labeling (∼64% of 488 TH+ neurons) of TH and CTb in the VTA, demonstrating the projection from VTA_DA_ neurons to the LH (**Figure 2B**).

**Figure 2.**
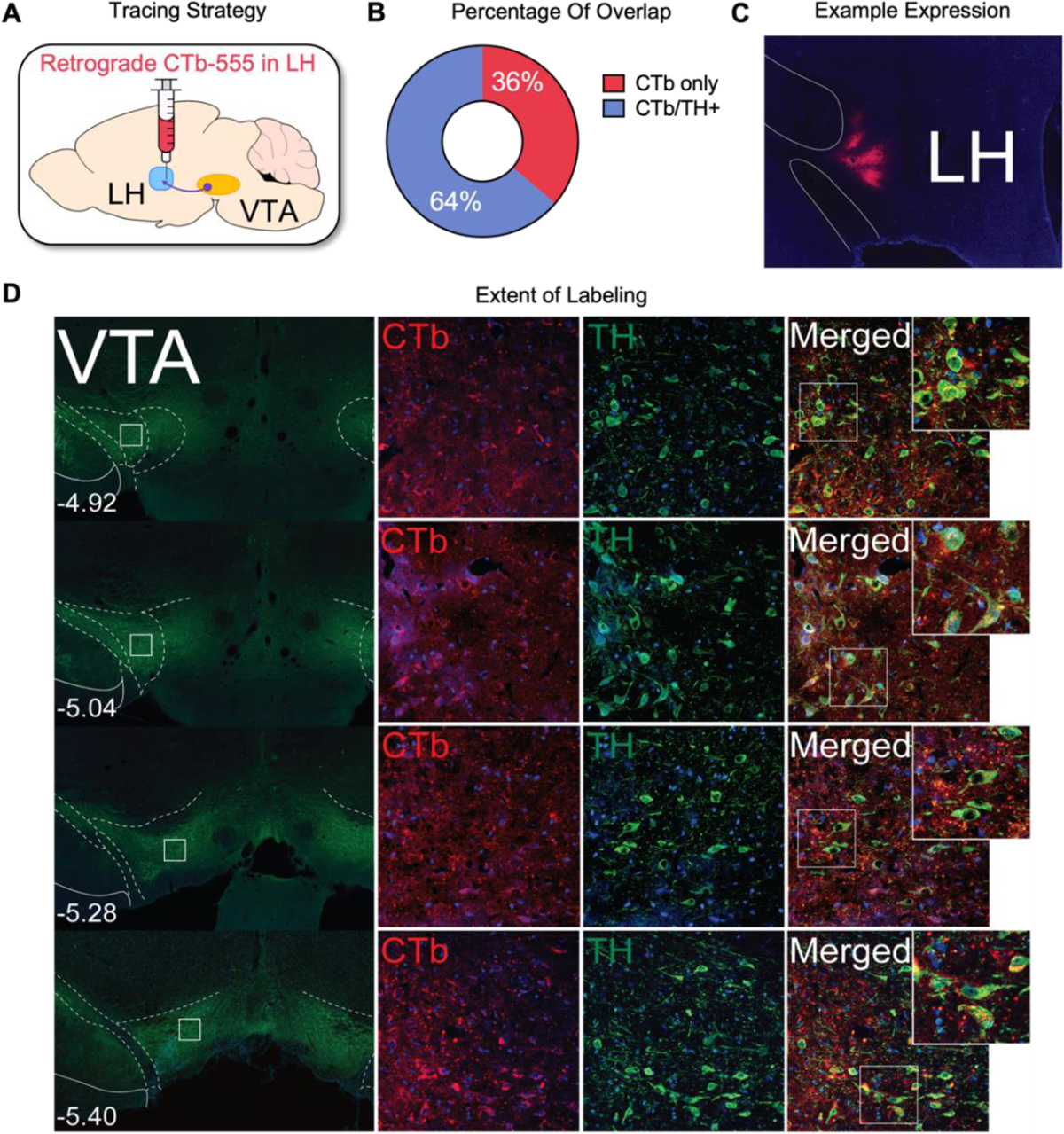
Revealing a novel dopaminergic projection to LH from VTA. (A) Schematic of retrograde tracing approach: rats were injected with cholera toxin subunit B (CTb-555) into LH. (B) Colocalization of CTb tracer and TH expression reveals ∼64% overlap in LH-projecting VTA cell bodies (*n*=377/488). (C) Example of CTb expression in LH. (D) Extent of CTb expression in VTA across the anterior/posterior plane relative to bregma, stained for CTb (red), TH+ (green), and overlap (merge) at 10x (left) and 20x (right, inset) magnification.

### VTA_DA_ projections to LH are necessary for model-based associations between cues and rewards

After verifying the existence of a VTA_DA_ projection to LH, we asked whether this circuit was necessary for the development of model-based associations in LH. We first bilaterally infused TH-Cre rats with a Cre-dependent NpHR virus (*n=*12) or eYFP control vector (*n*=10) into VTA and placed our optic fibers in LH (**Figure 3A-C**). This would allow us to selectively inhibit VTA_DA_ terminals in LH during learning when this pathway would be active under normal circumstances^35^. Following virus incubation, rats were food restricted and received Pavlovian conditioning (14 sessions; 12 presentations/session), where one 10-sec auditory cue leads to food reward (CS+), and another was without consequence (CS-; white noise or click; counterbalanced). During learning, green laser light was delivered at the time of reward following CS+ presentation (2.5-sec; 532nm; 16-18mW) using parameters that have been shown to suppress dopamine firing without causing a negative prediction error ^42, 69^. Specifically, these parameters do not produce extinction learning, which is seen with shorter bursts of inhibition that better mimic a negative prediction error ^30, 69^. We found that inhibition of VTA_DA_ terminals in LH reduced learning about the CS+ and not the CS-. This was supported by statistical analyses, revealing that both eYFP and NpHR groups elevated responding to CS+ relative to the CS-(**Figure 3D**; CS+ vs. CS-: *F*_(1,20)_ = 46.083, *p*<0.001; session: *F*_(6,120)_ = 5.582, *p*<0.001; group: *F*_(1,20)_ = 2.017, *p*=0.171; CS x session: *F*_(6,120)_ = 20.323, *p*<0.001; CS x group: *F*_(1,20)_ = 0.903, *p*=0.353; session x group: *F*_(6,120)_ = 1.571, *p*=0.161). However, there was a significant interaction between the groups and the rate at which they elevated their responding to the CS+ (CS x session x group: *F*_(6,120)_ = 3.788, *p*=0.002), which was most pronounced in the final session (simple main effect of group, CS+: *F*_(6,120)_ = 18.681, *p*=0.004), and without between-groups differences in CS-responding (simple main effect of group, CS-; *F*_(6,120)_ = 3.287, *p*=0.244). These results show that inhibition of VTA_DA_ terminals in LH impaired the ability of rats to learn about reward-predictive cues.

**Figure 3.**
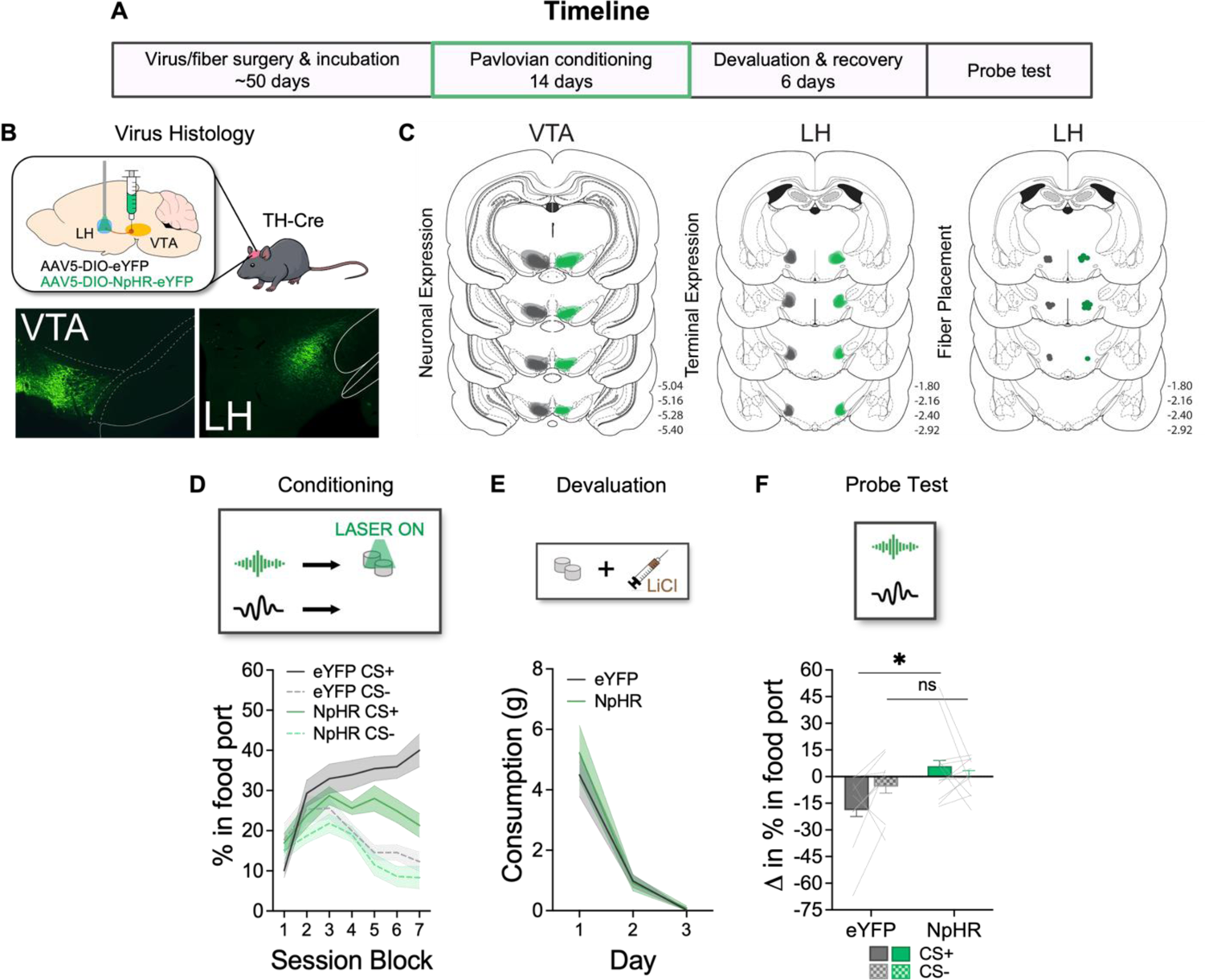
Inhibition of VTA_DA_ projections to LH reduces model-based learning about cues and rewards. (A) Experimental timeline. (B) Optogenetic approach: TH-Cre transgenic rats were bilaterally infused with a Cre-dependent AAV with inhibitory halorhodopsin (NpHR; *n*=12), or a control vector (eYFP; *n*=10) in VTA and implanted with optic fibers in LH to allow for the inhibition of VTA_DA_ terminals in LH. Below shows unilateral examples of bilateral virus expression in VTA_DA_ neurons (*left*) and axonal terminals in LH (*right*). (C) *Left*: Unilateral representation of bilateral cell body virus expression in VTA for the eYFP group (grey) and the NpHR group (green). *Middle:* Unilateral representation of bilateral axonal terminal expression in LH. *Right*: Dots indicate approximate location of fiber tips in LH. (D) Rats learned that a CS+ leads to reward and a CS-has no consequence. VTA_DA_ terminals in LH were inhibited during food delivery across learning, when a prediction error would occur. Inhibition of VTA_DA_ terminals in LH significantly reduced learning about the CS+. (E) Reward was then paired with injections of LiCl and both eYFP and NpHR groups reduced their consumption across LiCl pairings. (F) Rats received a probe test where the CS+ and CS-were presented without reward. Here, rats in the eYFP group reduced responding to CS+ after devaluation, while the NpHR group showed no difference. \**p* ≤ 0.05, mean (± SEM).

Although learning about the CS+ was reduced in the NpHR group of rats, responding was not completely abolished by VTA_DA_→LH inhibition. This afforded the opportunity to probe the nature of the learning that remained in these rats. To do so, we devalued the reward paired with CS+ by pairing the reward with injection of LiCl. Both eYFP and NpHR groups reduced their consumption of the reward with consecutive pairings of LiCl injections (**Figure 3E**; day: *F*_(2,40)_ = 55.124, *p*<0.001; group: *F*_(1,20)_ = 0.297, *p*=0.592; day x group: *F*_(2,40)_ = 0.394, *p*=0.677). Finally, rats were given a probe test to examine the effects of reinforcer devaluation on responding to the CS+. Given CS+ responding was reduced in the NpHR group relative to the eYFP group during learning, we compared the change in responding to the CSs before and after the devaluation procedure. Here, we found that the eYFP group reduced responding to the CS+ after devaluation, indicated by a negative change in responding to the CS+ but not the CS-. However, the NpHR group failed to show any change in responding to the CS+ (**Figure 3F**). This was supported by statistical analyses, which revealed no main effect of CS (CS+ vs. CS-: *F*_(1,20)_ = 0.595, *p*=0.449), but a significant main effect of group (eYFP vs. NpHR: *F*_(1,20)_ = 6.671, *p*=0.018) and a significant CS x group interaction (*F_(_*_1,20)_ = 3.827, *p*=0.033) owed to a significant difference in responding to the CS+ between the eYFP and NpHR groups (*F*_(1,20)_ = 7.646, *p*=0.012), and not the CS-*F*_(1,20)_ = 0.914, *p*=0.350). Importantly, there was no effect of trial (*F*_(5,100)_ = 1.077, *p*=0.378) or any interaction between trial and CS (*F*_(5,100)_ = 0.321, *p*=0.899), confirming an effect of devaluation and not extinction. This demonstrates that the *residual* learning to the CS+ was insensitive to reward devaluation, indicating what is learned in the VTA_DA_→LH circuit is model-based.

### Phasic stimulation of VTA_DA_ projections to LH is sufficient to drive model-based learning between cues and rewards

We found that inhibition of VTA_DA_ projections to LH attenuates model-based learning about cues and their specific rewards. However, given there was reduced responding in our NpHR experimental group, it was difficult to definitively say that there was a change in devaluation sensitivity. In order to address this, we next asked if stimulation of the VTA_DA_→LH pathway would be sufficient to drive model-based learning between cues and rewards (**Figure 4A**). To test this, we utilized the blocking procedure ^33, 38, 70, 71^, which allows us to test if we can biologically rescue associative learning by stimulation of the VTA_DA_ to LH pathway. TH-Cre rats were bilaterally infused with a Cre-dependent, excitatory channelrhodopsin (AAV5-Ef1a-DIO-hChR2(E123T/T159C)-eYFP; ChR2; *n*=14) in VTA and had an optic fiber placed over LH (**Figure 4B-C**). This allows us to stimulate VTA_DA_ terminals in LH. After virus incubation, rats were food restricted and began Pavlovian conditioning, where two 30-sec visual cues were paired with two distinct rewards (flash and steady lights, counterbalanced; 8 sessions; 8 presentations/session). We then introduced two novel 30-sec auditory cues (click and white noise, counterbalanced) presented in compound with the visual cues and followed by the same distinct rewards (4 sessions; 8 presentations/session). Normally, rats will not learn about the novel auditory cues because no new information can be attributed them as there is no change in reward identity or magnitude (i.e., blocking) ^70^. However, during one of the rewards, we stimulated VTA_DA_ terminals in LH as a prediction error by delivering blue light into LH (1-sec; 20Hz, 473nm, 14-16mW) ^41, 42, 72^, to examine whether we could facilitate learning about one of the auditory cues (“unblocked”) while the other cue would serve as a control is not learned about (“blocked”). As we and others have previously shown that light alone in eYFP controls does not unblock learning using these parameters ^30, 41, 42, 71^, we opted for a within-subjects blocking design where all rats had ChR2 infused into VTA_DA_ neurons which allowed us to compare responding to unblocked and blocked cues in each rat. All rats elevated responding above baseline to the visual stimuli in initial conditioning sessions, and this was unaffected by introduction of the auditory cues and stimulation of the VTA_DA_ to LH circuit (**Figure 4D-E**; time period: *F*_(2,26)_ = 49.022, *p*<0.001; session: *F*_(5,65)_ = 3.500, *p*=0.007; time period x session: *F*_(10,130)_ = 7.938, *p*<0.001; blocked vs. baseline: *F*_(10,130)_ = 3.442, *p*<0.001; unblocked vs. baseline: *F*_(10,130)_ = 3.156, *p*<0.001; blocked vs. unblocked: *F*_(10,130)_ = 0.286, *p*=0.550). Rats then received a probe test in which each of the auditory cues were presented without rewards or stimulation. We found that rats made greater responses to the unblocked cue than the blocked cue (**Figure 4F**; *F*_(1,13)_ = 4.28, *p*=0.030) demonstrating that phasic stimulation of the VTA_DA_ terminals in LH successfully facilitated learning about a reward-paired cue.

**Figure 4.**
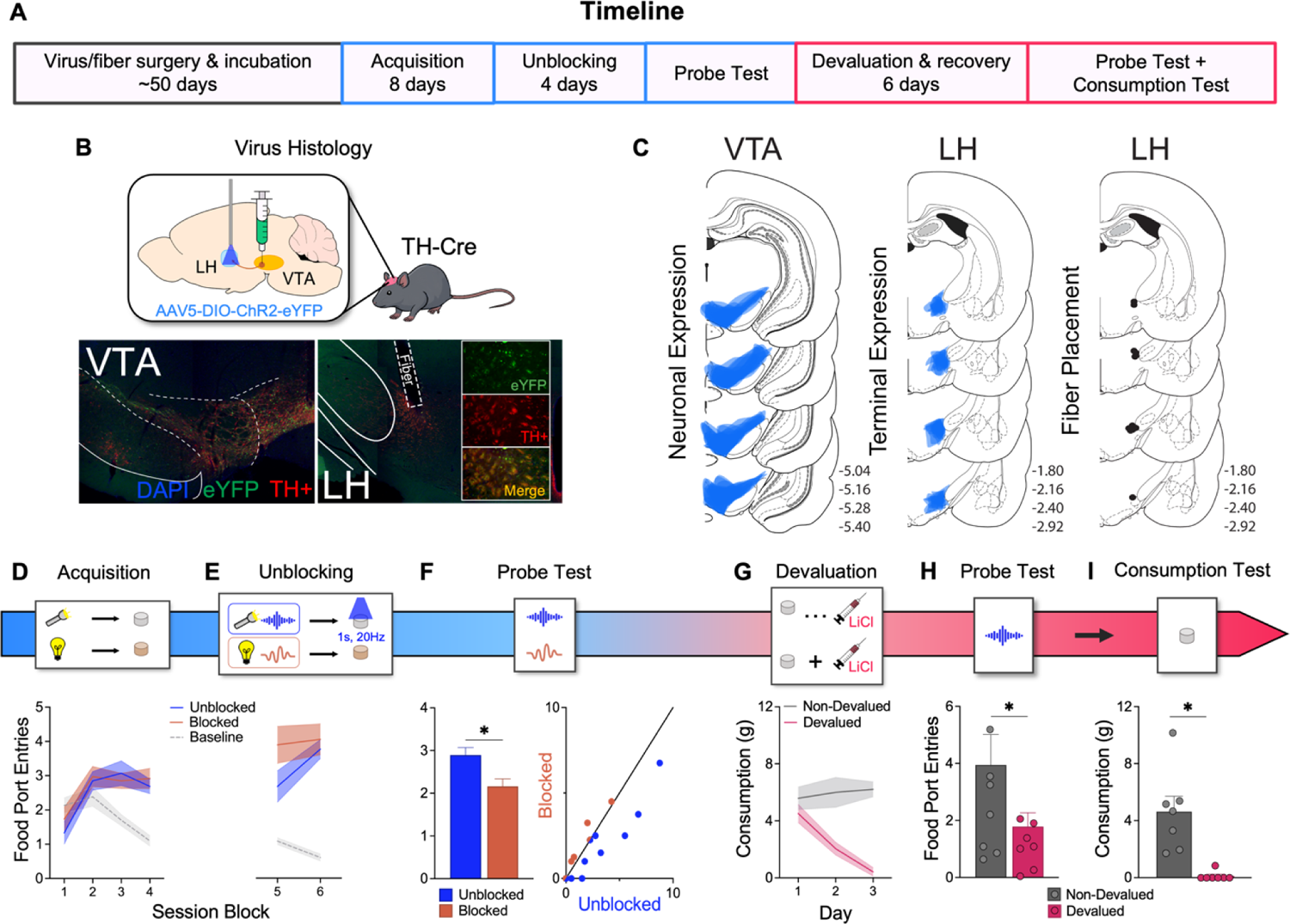
Phasic stimulation of the VTA_DA_→LH pathway facilitates model-based learning for cues and rewards. (A) Experimental timeline. (B) Optogenetic approach: TH-Cre transgenic rats were bilaterally infused with a Cre-dependent AAV with channelrhodopsin (*n*=14) in VTA and implanted with optic fibers in LH. *Middle:* unilateral examples of bilateral virus expression in the cell bodies of dopamine neurons in VTA and *Bottom*: in axonal terminals in LH. Slices were stained with DAPI (blue) and TH (red) for immunohistochemical verification of co-localization of virus expression (tagged with eYFP, green) and TH. (C) *Left*: Unilateral representation of bilateral virus expression in VTA. *Middle:* Unilateral representation of bilateral terminal expression in LH. *Right*: Black dots indicate approximate location of fiber tips in LH. (D) Rats first learned an association between two visual cues and two distinct rewards. Responding is represented as the number of entries made into the food port during cue presentation across session blocks (2 daily sessions per block). (E) Novel auditory cues were presented in compound with the visual cues and led to the same rewards. Blue light (20Hz, 473nm, 14-16mW) was delivered concurrently with one of the rewards. (F) Rats were then given a probe test for the auditory cues. Here, rats showed significantly more responding to the cue paired with stimulation of VTA_DA_ terminals in LH (i.e., unblocked cue). Individual responding is represented as dots on the scatterplot, with color indicating a preference for blocked (orange) or unblocked (blue) cue. To the extent that responding is equivalent to these cues, dots will congregate on the diagonal. (G) Rats underwent a reward devaluation procedure where the reward associated with the unblocked cue was paired with injections of LiCl. The devalued group, but not the non-devalued group, reduced consumption of the reward across LiCl injections. (H) Rats received a final probe test to examine the devaluation-sensitivity of responding to the unblocked cue. Here, the devalued group made significantly fewer responses to the cue. (I) Rats also underwent a consumption test to confirm that devaluation procedures were effective. The devalued group consumed less of the reward than the non-devalued group. \**p* ≤ 0.05, mean (± SEM).

Finally, we probed the content of learning supported by stimulation of the VTA_DA_→LH pathway. To investigate this, we employed a devaluation procedure in which half of the rats received pairings of LiCl injections with consumption of the reward associated with the unblocked cue (“devalued”) and the other half would experience LiCl injections separate from reward consumption (“non-devalued”). The non-devalued group maintained consumption of the reward across devaluation days, while the devalued group reduced consumption, demonstrating development of a conditioned taste aversion (**Figure 4G**; day: *F*_(2,24)_ = 6.064, *p*=0.007; group: *F*_(1,12)_ = 22.561, *p*<0.001; day x group: *F*_(2,24)_ = 11.412, *p*<0.001). Rats then received a probe test where the unblocked cue was presented alone without reward. Here, the devalued group exhibited a reduced level of responding to the cue relative to the non-devalued group (**Figure 4H**; Welch’s test: *F*_(1,13)_ = 3.404, *p*=0.050). This suggested that learning about the unblocked cue was model-based. Further, a consumption test for the devalued reward was conducted immediately following the probe test, which confirmed devaluation procedures were successful (**Figure 4I**; Welch’s test; *F*_(1,13)_ = 17.247, *p*=0.003). In accordance with the data from inhibition of this circuit (**Figure 3**), these data suggest that the VTA_DA_→LH pathway functions to support learning of model-based cue-reward associations.

### Methamphetamine self-administration enhances use of model-based associations to drive behavior and sensitizes the hypothalamic-midbrain circuit

Here, we have shown that the LH encodes model-based associations and that this is facilitated by input from VTA_DA_ neurons. Interestingly, prior research has implicated a strengthening of hypothalamic circuits in the neural changes that occur with exposure to drugs of abuse ^11–17^. Specifically, it has been shown that the LH undergoes robust gene expression changes for pre- and postsynaptic proteins involved in neurotransmission following cocaine self-administration ^17^ and that increased Fos activity of neurons in this region after exposure to drugs of abuse is associated with addiction-like phenotypes ^11, 15, 16^. LH neurons are also activated during context-induced relapse of drug- and alcohol-seeking ^15, 20, 21, 73, 74^. Further, exposure to substances such as cocaine or morphine increase Fos expression of LH neurons projecting to VTA in response to drug-predictive cues and contexts ^12, 13^. If the neural changes that occur in this circuit are relevant for changes in learning and behavior that occur with drugs of abuse, this would lead to the prediction that drug exposure would enhance the use of model-based associations between cues and rewards to influence decision-making. This is counterintuitive as we generally conceptualize reinforcement learning following drug abuse to bias towards model-free systems. For example, a classical interpretation of decision-making deficits in drug addiction is that they respond in a habitual manner that is devoid of consideration of outcome (i.e., stimulus-response associations) ^3–10^. However, most of the supporting evidence for this has involved the use of instrumental procedures (i.e., action-dependent) and not Pavlovian procedures (i.e., action-independent) like those used in the present study. To examine how addictive substances like methamphetamine would impact the use of model-based Pavlovian associations between cues and rewards, we tested how prior self-administration of methamphetamine affects performance in the specific Pavlovian-to-instrumental transfer task. This task would allow us to assess: 1) how much control Pavlovian cues have over behavior after drug exposure, and 2) whether the nature of cue control is related to the specific outcome it predicts (i.e., model-based) or generally invigorates rewarded responses (i.e., model-free).

Experimentally-naïve rats first underwent surgery for the implantation of intravenous catheters in their jugular vein. Rats were then food restricted and trained to self-administer either grain pellets (control, *n*=8) or methamphetamine infusions [methamphetamine, *n*=7; (**Figure 5A**)]. Self-administration sessions comprised of 3-hour sessions across a 14-day protocol, beginning with rats pressing the active lever once for rewards (fixed-ratio 1; FR1) and escalating to FR3 and then FR5 reinforcement schedules ^75–77^. In the control group, pressing the active lever resulted in delivery of two 45-mg grain pellets (Test Diet, MA). In the methamphetamine group, pressing the active lever resulted in a 0.1 mg/kg intravenous methamphetamine infusion. For both groups, pressing the inactive lever had no programmed consequences. The control group increased lever-pressing on the active lever across time relative to the inactive lever (**Figure 5B (left)**; lever: *F*_(1,7)_ = 256.163, *p*<0.001; session: *F*_(13,91)_ = 54.412, *p*<0.001; lever x session: *F*_(13,91)_ = 65.966, *p*<0.001), which was also reflected in them earning more pellets (g/kg) across time (**Figure 5B (right)**; session: *F*_(13,91)_ = 7.699, *p*<0.001). Similarly, the methamphetamine group also increased responding on the active lever relative to the inactive lever (**Figure 5C (left)**; lever: *F*_(1,6)_ = 16.641, *p*=0.007; session: *F*_(13,78)_ = 23.628, *p*<0.001; lever x session: *F*_(13,78)_ = 9.241, *p*<0.001). This was also reflected in escalation of their methamphetamine intake across time (mg/kg) (**Figure 5C (right)**; session: *F*_(13,78)_ = 6.119, *p*<0.001).

**Figure 5.**
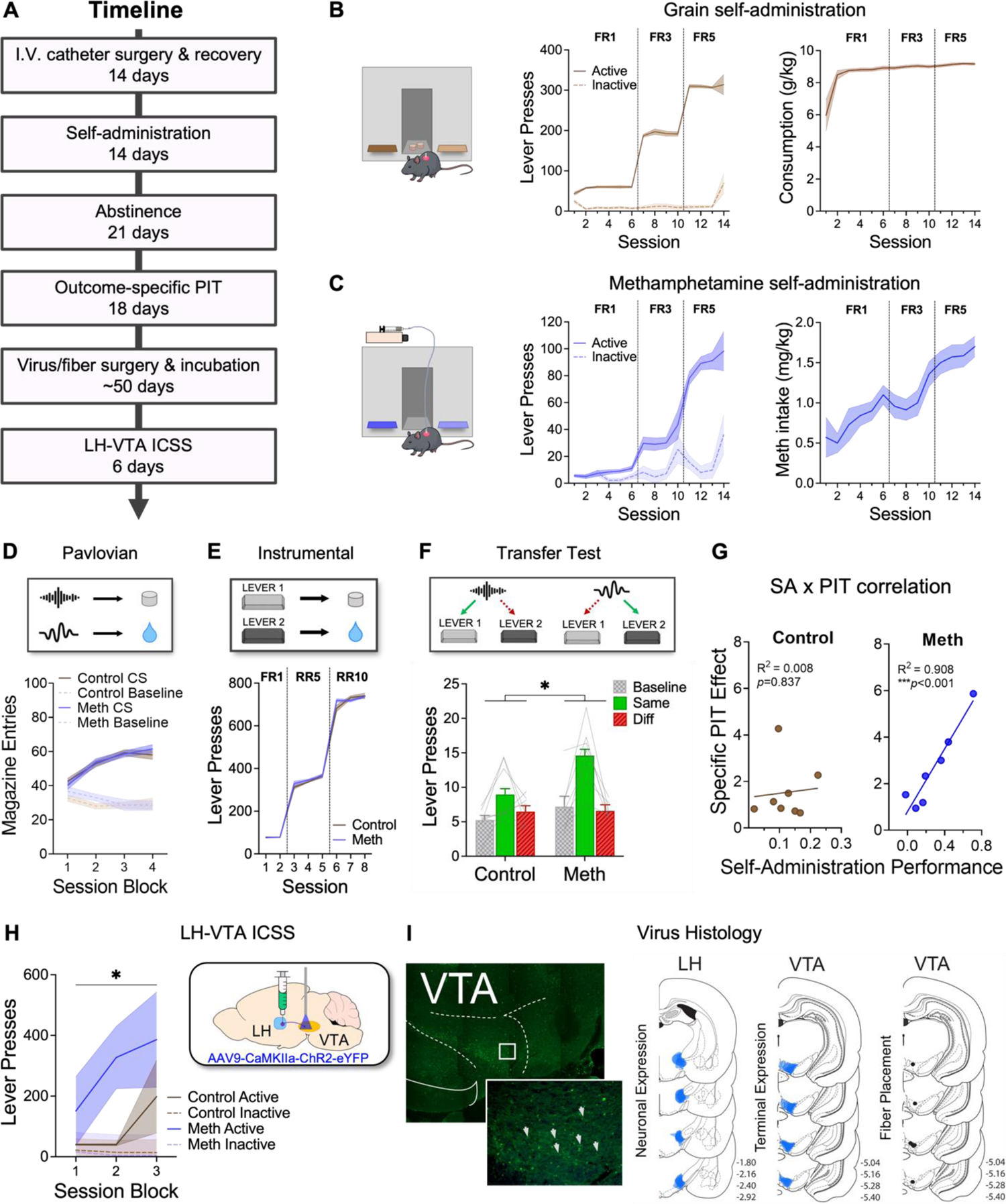
A history of methamphetamine self-administration enhances specific PIT and strengthens the LH→VTA pathway. (A) Experimental timeline. (B) Rats in the control group first learned to press a lever for grain. These rats increased responding on the active lever (*left*), resulting in an increase in pellets earned (*right*). (C) Rats in the methamphetamine group learned to press a lever for 0.1 mg/kg infusions of methamphetamine. Lever-pressing for drug escalated across time (*left*) as well as their methamphetamine intake across time (*right*). (D) Following abstinence, all rats then received Pavlovian conditioning, increasing food port entries across sessions. (E) Next, rats learned the instrumental contingencies and increased responding for the two distinct rewards across time. (F) Finally, rats underwent the critical PIT test. Here, the methamphetamine group showed an enhanced specific PIT effect. (G) There was no correlation between grain self-administration and the magnitude of specific PIT (*left*), however, a strong positive correlation was present between methamphetamine self-administration and the outcome-specific PIT effect (*right*). (H) We tested how much rats were willing to earn stimulation of the LH→VTA pathway (intracranial self-stimulation; ICSS). Here, the methamphetamine group showed significantly greater ICSS, implicating sensitization of the LH-VTA pathway. (I) Unilateral example of bilateral virus expression in the axonal terminals in VTA (*left*), and schematics of virus expression in the LH and VTA with fiber placements in VTA (*right*). \**p* ≤ 0.05, mean (± SEM).

Three weeks after self-administration, both groups were trained on a specific Pavlovian-to-Instrumental Transfer (PIT) paradigm in a new context. First, rats learned two Pavlovian contingencies ^78^. Here, a 2-min auditory cue (click and white noise, counterbalanced; 8 sessions, 8 presentations/session) predicted one of two distinct rewards [45-mg sucrose pellets (Test Diet, MA) and 15% maltodextrin solution (Earthborn Elements, OR); counterbalanced]. Both control and methamphetamine groups readily acquired the conditioned food-port entry during the cues without between-groups differences (**Figure 5D**; CS vs. pre-CS: *F*_(1,13)_ = 137.677, *p*<0.001; session: *F*_(3,39)_ = 4.406, *p*=0.009; group: *F*_(1,13)_ = 0.053, *p*=0.822; CS vs. pre-CS x group: *F*_(1,13)_ = 0.195, *p*=0.666; session x group: *F*_(3,39)_ = 0.244, *p*=0.865; CS vs. pre-CS x session: *F*_(3,39)_ = 17.374, *p*<0.001; CS vs. pre-CS x session x group: *F*_(3,39)_ = 0.778, *p*=0.513). Rats were then trained to press two different levers to receive the two outcomes (e.g., left lever→sucrose, right lever→maltodextrin; counterbalanced) across increasing random-ratio (RR) reinforcement schedules (FR1, RR5, RR10; 8 sessions). Both groups readily acquired instrumental contingencies without between-group differences (**Figure 5E**; session: *F*_(7,91)_ = 477.738, *p*<0.001; group: *F*_(1,13)_ = 0.053, *p*=0.821; session x group: *F*_(7,91)_ = 0.386, *p*=0.908).

Finally, rats were given the PIT test in which both levers were available and each of the auditory cues were presented. Importantly, no rewards were delivered so that we could test for the representation evoked by the cues without reward feedback. Specific PIT is observed when greater responses are made on the lever predicting the same outcome as the presented cue (“Same”), relative to the lever leading to the alternative outcome (“Diff”). This illustrates the use of model-based associative information to drive decision-making ^79–82^. We found that while both groups responded more on the same lever relative to the different lever and above baseline responding prior to cue onset (**Figure 5F**; lever: *F*_(2,26)_ = 15.694, *p*<0.001), the methamphetamine group showed overall heightened responding compared to controls (group: *F*_(1,13)_ = 6.119, *p*=0.028), driven by a marked increase in the magnitude of the difference between responding on the same and different levers (lever x group: *F*_(2,26)_ = 3.229, *p*=0.028). This was due to a significant group difference in responding on the same lever (*F*_(2,26)_ = 5.670, *p*=0.024) without differences in responding on the different lever (*F*_(2,26)_ = 0.103, *p*=0.913) nor in baseline responding (F_(2,26)_ = 1.977, *p*=0.229), indicating an enhancement of the specific PIT effect. Indeed, while the methamphetamine group showed a significant difference in responding on the same vs. diff levers when analyzed independently (simple main effect: *F*_(2,26)_ = 8.036, *p*<0.001), our control group did not (simple main effect: *F*_(2,26)_ = 2.469, *p*=0.168). However, both groups displayed elevated responding for the same lever above baseline (same vs. baseline, methamphetamine: F_(2,26)_ = 7.399, *p*<0.001; control: F_(2,26)_ = 1.509, *p*=0.029) without differences for the different lever relative to baseline (diff vs. baseline, methamphetamine: F_(2,26)_ = 0.637, *p*=0.666; control: F_(2,26)_ = 1.237, *p*=0.376). Moreover, we found a strong positive correlation between self-administration and the magnitude of specific PIT in the methamphetamine group (**Figure 5G (right)**; R^2^ = 0.908, *p*<0.001) but not the control group (**Figure 5G (left)**; R^2^ = 0.008, *p*=0.837). These data show that methamphetamine self-administration produced an increase in the influence of model-based cue-reward associations over decision-making. An enhancement in specific PIT was further supported by a replication of this effect in rats receiving experimenter-administered injections of methamphetamine rather than self-administration of methamphetamine (**Supplemental Figure 1**). This confirmed that it was not the *lever pressing* for methamphetamine (or grain pellets) that produced the changes in the specific PIT effect. Rather, methamphetamine itself was sufficient to drive the changes in reward learning.

Next, we asked whether changes in our novel LH-VTA circuit could result from methamphetamine self-administration that may be related to enhancements in specific PIT. To test this, we compared how much our control and methamphetamine groups would show intracranial self-stimulation (ICSS) for the LH→VTA pathway, where ICSS is driven by GABAergic input from LH to VTA ^83^. A subset of rats from our control and drug groups underwent surgeries to bilaterally infuse channelrhodopsin (AAV9-CaMKIIa-hChR2(H134R)-eYFP) in LH and implant an optic fiber placed over VTA, allowing for stimulation of LH terminal projections in VTA (**Figure 5I**). Rats were then given ICSS sessions in a new context (**Figure 5H**; 30-min sessions, 6 sessions). Here, rats could press an active lever that delivered light-mediated stimulation of LH terminals in VTA (2-sec; 20Hz, 473nm, 14-16mW) or an inactive lever which had no programmed consequences. We found that the methamphetamine group showed significantly faster acquisition of ICSS relative to the control group, illustrated by steeper increases in active lever presses across time (session x lever x group: *F*_(2,8)_ = 6.806, *p*=0.019; simple effect of lever during the 2^nd^ block: *F*_(2,8)_ = 1.735, *p*=0.044). This demonstrates that the input from LH to VTA is enhanced in our rats experiencing methamphetamine self-administration. Interestingly, we also found evidence suggesting a strengthening of the VTA→LH pathway with methamphetamine exposure. Specifically, we infused channelrhodopsin into VTA and optic fibers into the LH of rats that received experimenter-administered methamphetamine and allowed them to undergo the ICSS procedure for stimulation of the VTA→LH pathway. We found that rats with a history of methamphetamine displayed enhanced ICSS relative to controls (**Supplemental Figure 2**), similarly to rats afforded the opportunity to press a lever for stimulation of the LH→VTA pathway. Altogether, these data implicate a strengthening of the LH-VTA circuits following a history of methamphetamine self-administration that may be contribute to the enhancements in LH-dependent model-based processes.

## Discussion

In the present study, we set out to understand the nature of learning that underlies LH function, the wider circuity that supports this function, and whether drug exposure enhances learning dependent on LH circuity. Our first study revealed that LH_GABA_ neurons are necessary for the use of model-based associations to guide behavior. Specifically, we found that rats without LH_GABA_ neuronal activity were unable to represent the current value of future rewards that facilitates flexible behavior. This extends our previous work demonstrating that LH_GABA_ neurons are needed to acquire and express learning about cues and rewards ^19^, revealing that the nature of this learning involves the development of model-based associations comprised of representations of cues and their specific predicted rewards. In the context of our work demonstrating that LH_GABA_ neurons bias learning toward cues proximal to the reward ^18^, we can now view this bias toward proximal cues exhibited by the LH as a model-based phenomenon. This has interesting implications for the theoretical frameworks of model-based learning ^1^, which do not distinguish between neural substrates that are responsible for the distal and proximal steps in the model-based cognitive map. Thus, we would argue that LH_GABA_ neurons are preferentially involved in biasing learning towards cues proximal to rewards in a model-based manner, revealing a neural dissociation in the proximal and distal features of model-based associations.

Next, we identified a novel projection from VTA dopamine neurons to the LH, which we revealed is both necessary and sufficient to support model-based learning in LH. This is notable because it is one of the first implications of a specific and direct circuit comprising dopamine neurons that contributes to model-based learning ^84, 85^. Previous anatomical studies have suggested that a projection from VTA dopamine neurons to LH may exist ^48, 49, 67, 68^. Here, we used a retrograde tracing strategy to confirm and quantify this dopaminergic projection from VTA to the LH. We then demonstrated that inhibition of VTA dopamine terminals in LH during cue-reward learning significantly reduced the ability of rats to use the cue to predict upcoming rewards. Further, subsequent devaluation procedures revealed that the learning that was still intact after inhibition of the VTA dopamine to LH pathway lacked a representation of the reward, demonstrating that the model-based component of that learning was abolished. We then showed that stimulation of VTA dopamine terminals in LH was also sufficient to drive new learning about cues and rewards, where the learning supported by stimulation of this pathway was again revealed to be model-based in nature. It is important to note here that stimulating the terminals in LH from VTA dopamine could have impacted the fibers of passage on route to nucleus accumbens. While this is possible, we do not think it could drive our results because we found the same result with inhibition of these terminals, which is unlikely to impact processes distant from the terminals, and we separately confirmed a large proportion of VTA dopamine neurons project directly to LH. Importantly, using the temporal specificity of optogenetics, we implicated this circuit in a manner consistent with the endogenous function of VTA dopamine neurons. That is, our manipulations were precisely timed to closely reflect the endogenous firing patterns of VTA dopamine neurons ^35, 86^. We inhibited VTA dopamine terminals in LH during reward receipt across a period of time that would suppress a potential prediction error without producing a negative prediction error ^69^. Similarly, we stimulated VTA dopamine terminals in LH during reward receipt using temporal parameters that follow the endogenous firing of dopamine neurons ^41, 42, 87^. Thus, our manipulations approach physiological conditions of the circuit. These data are consistent with modern accounts of dopamine prediction errors in supporting associative learning beyond scalar, model-free value ^36–46^ and reveal a novel hypothalamic-midbrain circuit that underlies model-based learning about cues and rewards.

Given the knowledge that the input from LH to the VTA is strengthened following psychostimulant exposure ^12, 13^, we were interested in testing whether self-administration of methamphetamine would enhance model-based learning about cues and rewards, which we have shown in the present experiments is dependent on this hypothalamic-midbrain circuit. To test this, we first allowed rats to self-administer methamphetamine and then learn associations between two cues and two rewards. After instrumental training for two actions and these same rewards, we then examined the extent to which the cues would invigorate actions directed towards the paired rewards. Prior research has shown that drugs of abuse increase the impact of cues in invigorating responding directed to rewards in both rats and humans ^27, 88–97^ and our use of the specific Pavlovian-to-Instrumental transfer procedure could reveal the nature of this enhancement. Indeed, we found that the cues exerted heightened control over behavior in a manner that reflected behavior directed towards specific rewards after drug exposure, indicating a model-based process ^1, 79^. These enhancements in model-based decision-making as a result of prior drug exposure is surprising as it counters the canonical habit, or model-free, theory of addiction ^3-10^. That is, substance use disorder is often conceptualized as promoting habitual behavior directed towards drug rewards, which explicitly lacks a representation of the specific reward. Indeed, this could be in part because persistent drug seeking alters the perception of the instrumental cost to obtain rewards following drug exposure ^76, 98^. Our data suggest that this development of habits following psychostimulants like amphetamines ^6, 99, 100^ may be specific to instrumental responding, and that the influence of Pavlovian cues over drug seeking and drug taking is in fact model-based. A more encompassing perspective on the reinforcement learning mechanisms underlying substance use disorder could integrate the model-based influence that drug-paired cues have over habitual instrumental behaviors directed towards drug taking. This would facilitate a better understanding of the complexities of drug taking in naturalistic environments that comprise both instrumental and Pavlovian components. Such a perspective could reconcile contradictions in the literature as to whether drug seeking is the result of habitual or goal-directed processes ^8, 101, 102^, demonstrating that it depends on whether the influence is based on instrumental or Pavlovian processes.

There were also two notable features of the enhancement of model-based behavior seen following methamphetamine exposure. The first is that we found an enhancement in the specific Pavlovian-to-Instrumental transfer effect regardless of how the exposure to methamphetamine occurred. Specifically, we demonstrated that both self- and experimenter-administered methamphetamine enhanced the specific Pavlovian-to-instrumental transfer effect. This is interesting given controversy as to whether it is the act of taking drugs that changes the neural circuits involved in reward learning that can influence future drug seeking ^103^, or whether the drug itself is sufficient to produce neural changes underlying reward learning ^6^. Here, we found evidence that methamphetamine itself is sufficient to produce the heightened control that reward cues have over behavior. The second point to note is that we found the increase in model-based control of cues over behavior was also accompanied by a strengthening of this hypothalamic-midbrain pathway. That is, we demonstrated that rats previously receiving methamphetamine self-administration would press more for stimulation of LH input to the VTA, which indicates a strengthening of the GABAergic input to VTA that mediates the reinforcing properties within this circuit ^83^. We further showed evidence to suggest that the input from VTA to the LH is also strengthened with drug exposure. Indeed, rats given experimenter-administered methamphetamine also pressed more to receive stimulation of VTA input to the LH. These findings are in line with other accounts showing neuroplasticity and increased activation of hypothalamic circuits following exposure to drugs of abuse ^11–13, 15–17, 73, 74^, and suggest for the first time that this novel input from VTA neurons to the LH is also strengthened following psychostimulant exposure. Of course, future research is necessary to determine the way in which these circuits are changed following drug exposure. For example, it could be the case that the depletion of dopamine transporters following methamphetamine use ^104–107^ means drug-exposed subjects require more stimulation to experience the same level of reinforcement as in drug-naïve subjects. While this would be inconsistent with other research showing increased activity in these circuits ^11–13, 15–17, 73, 74^, it warrants further consideration. Nevertheless, these results suggest that methamphetamine produces enhancements in the model-based control that cues have over behavior and that this is accompanied by changes in the bidirectional LH-VTA circuit.

Combined with our prior data ^18, 19^, we now understand that the hypothalamic-midbrain circuit contributes to model-based learning about cues and rewards in a manner that biases learning about cues most proximal to reward. We would argue that the strengthening of the LH-VTA circuit following drug exposure enhances the bias in learning and behavior directed towards reward-paired cues, which increases the control that these cues have over decision-making relative to other information in the environment that may not be directly reward relevant. This mirrors the pattern of reinforcement learning changes seen in humans with drug addiction, and rodent models of the disorder ^27, 75, 88, 90–95^. This reveals the LH-VTA circuit as a critical node in the reinforcement learning changes seen drug addiction, consistent with data implicating the LH in cue-induced reinstatement ^12, 14, 15, 20, 21, 73, 74, 108^. Beyond this, these data suggest targeting LH circuits could not only reduce the impact of drug cues on behavior, but also re-establish an appropriate balance in learning about other information in the environment.

Future research is needed to understand how the LH-VTA circuit integrates with the wider dopaminergic network. For example, we have previously hypothesized that the LH-VTA circuit forms a wider with the basolateral amygdala ^109^. Here, we argue that the basolateral amygdala provides the LH with sensory-specific information about motivationally significant events relevant to the current circumstance ^109^. This then allows LH to influence VTA dopamine signaling and bias learning towards information most relevant to current motivational states and goals ^19, 84, 109^. In contrast, given evidence that LH actively opposes learning about model-based information not directly related to rewards ^18^, it is also likely that the LH-VTA circuit acts to reduce the impact of other dopamine circuits in achieving their learning goals. For example, we have shown that inhibition of orbitofrontal circuit produces a dissociable effect from the LH on learning about distal model-based associations ^110^. This reveals a tension between the LH-VTA circuit and those comprising orbitofrontal cortex, which likely also involve input from VTA dopamine neurons ^85^. Altogether, this research begins to paint a picture of a complex and dynamic dopamine system, acting to establish a balance between many forms of learning. When this balance is impacted by external forces, like drug of abuse, it can skew this system towards particular forms of learning that produce changes in behavior characteristic of a pathological physical and psychological state.

## Materials and Methods

### Subjects

A total of 93 Long-Evans rats were used across all behavioral experiments in this study. For LH_GABA_ inhibition, 23 transgenic rats (10 female, 13 male) expressing Cre-recombinase under the control of the glutamate decarboxylase-1 (GAD) promoter were used (RRRC#751; Rat Resource and Research Center, MO). Optogenetic manipulations of the VTA_DA_→LH pathway used 36 different transgenic rats (14 female, 24 male) expressing Cre-recombinase under the control of tyrosine hydroxylase (TH) promoter (RRRC#659; Rat Resource and Research Center, MO). Pathway validation using immunohistochemical techniques used 4 non-transgenic, wild-type male rats (Charles River, MA). Experiments involving methamphetamine experience used 30 non-transgenic, wild-type rats (16 female, 14 male; Charles River, MA). Rats were randomly assigned to groups and matched for age and sex. Optogenetics experiments had rats maintained on a 12-hr light-dark cycle where all behavioral procedures were conducted during the light cycle. Methamphetamine experiments had rats on a 12-hr reverse light-dark cycle where training and testing were conducted during the early portion of the dark cycle. Prior to all training and testing procedures, rats were food restricted to ∼85% of their free-feeding body weight and maintained for the duration of the studies.

### Surgeries

#### Virus infusions and optic fiber implantation

General surgical procedures have been described elsewhere ^19, 42^. All surgical coordinates are relative to bregma. Rats were given 4-6 weeks to recover from surgical procedures and to allow for sufficient time for the virus to incubate in cell bodies and axonal projections. To optogenetically inhibit LH_GABA_ neurons, GAD-Cre rats were bilaterally infused with 1.0 µL of Cre-dependent adenoassociated virus carrying either inhibitory halorhodopsin (AAV5-Ef1a-DIO-eNpHR3.0-eYFP) or control virus without opsin (AAV5-Ef1a-DIO-eYFP) into LH (AP: −2.4 mm; ML: ±3.5 mm; DV: −9.0 (males) or −8.4 (females); angled at 10° towards midline). Optic fibers were also bilaterally implanted into LH (AP: −2.4 mm; ML: ±3.5 mm; DV: −8.5 (males) or −7.9 (females); angled at 10° towards midline). To optogenetically inhibit VTA dopamine terminals in LH, TH-Cre rats received bilateral infusions of 2.0 µL of either Cre-dependent halorhodopsin or control AAV into VTA [AP: −5.3 mm; ML: ±0.7 mm; DV: −7.0 and −8.2 mm (males) or −6.5 and −7.7 mm (females)]. Optic fibers were placed bilaterally over LH. Similar virus and fiber approaches were used for stimulation of the VTA_DA_→LH pathway, with the exception that TH-Cre rats were infused instead with 2.0 µL per hemisphere of Cre-dependent, excitatory channelrhodopsin (AAV5-Ef1a-DIO-hChR2(E123T/T159C)-eYFP), and fiber placed bilaterally over the LH. To establish intracranial self-stimulation of the LH→VTA pathway, rats received 1.2 µL bilateral infusions of CaMKIIa-driven channelrhodopsin (AAV9-CaMKIIa-hChR2(H134R)-eYFP) into LH (see virus coordinates for LH) and optic fiber implants over VTA (AP: −5.3 mm; ML: ±2.61 mm; DV: −9.0 (males) or −8.4 mm (females); angled at 15° towards midline). To allow for stimulation of the VTA→LH pathway, rats received 2.0 µL bilateral infusions of CaMKIIa-driven channelrhodopsin (AAV9-CaMKIIa-hChR2(H134R)-eYFP) into VTA with fibers placed bilaterally over LH.

#### Retrograde tracing

Rats were bilaterally infused with 0.6 µL per hemisphere of retrograde tracer, *Cholera Toxin Subunit B* (Thermo Fisher Scientific, MA), fluorescing at 555nm, into the LH using the same viral coordinates for this region as described earlier. Rats were allotted 6 days for tracer incubation before they were perfused for whole brain collection and immunohistochemical validation.

#### Intravenous catheterization

All rats in the self-administration experiment first received surgery to implant a homemade intravenous (I.V.) catheter into its jugular vein. Catheters consisted of a 14-cm length of Silastic tubing (I.D.: 0.012 in., O.D.: 0.025 in.) attached to a 22-gauge guide cannula with the distal end bent to a 90° angle embedded in dental acrylic anchored with a 2-cm square mesh. Rats were anesthetized with vaporized isoflurane gas and maintained at 2-3% vapor for the duration of the surgery. Rats received one 2-cm, lateral incision on its back between the shoulder blades and one 1-cm, vertical incision on its neck lateral to the midline. The base of the catheter was inserted subcutaneously through the back incision and the catheter tubing line exited through the neck incision. The right jugular vein was isolated and punctured with a 22-gauge needle before the catheter tubing was inserted into the vein and secured to surrounding muscle with silk sutures. Incisions were closed with surgical staples and vicryl sutures which were removed 1-week post-operation. The guide cannula of the catheter base was sealed with a small plastic cap and metal cover cap to help protect the catheter from debris. Catheters were flushed daily with 0.1 mL of saline and 0.2 mL heparinized saline containing enrofloxacin antibiotic (Baytril). Following catheterization, animals were randomly placed into either methamphetamine or control groups, allocated by sex and weight. Each group comprised of 8 rats (4 male and 4 female). Rats were given 1 week to recover from surgery before beginning self-administration procedures.

### Immunohistochemistry

Rats underwent transcardial perfusions, first using 1X phosphate buffered saline (PBS) solution then ice-cold 4% paraformaldehyde (PFA) solution made up in 1X PBS. Whole brains were collected and first stored in 4% PFA overnight at 20°C before transferring into 30% sucrose made up in 1X PBS. Brains were then sectioned into 20-40 µm slices using a cryostat and stored in PBS at 20°C. General procedures for immunohistochemistry are described elsewhere ^42, 111^. eYFP fluorescence was used to confirm TH+ expression in VTA cell bodies using a rabbit anti-TH antibody (1:1000; Millipore Sigma, Burlington, MA) as the primary antibody. A goat anti-rabbit IgG Alexa Fluor^TM^ 594 conjugate (1:500; Invitrogen, Waltham, MA) was used as the secondary antibody to stain for CTb tracer in VTA cell bodies. Slices were either washed using DAPI (4’,6-Diamidino-2-Phenylindole, Dihydrochloride; 1:10000; Thermo Fisher Scientific, MA) made up in di H_2_O to stain for nuclei before being mounted or cover-slipped with ProLong Gold mounting medium with DAPI (Fisher Scientific, MA) on slides for imaging with a Zeiss confocal microscope (Zeiss, Oberkochen, Germany) with 10x and 20x objectives.

### Quantification of neurons

Tissue from rats used for retrograde tracing (*n*=4) was imaged following immunohistochemical processing under a 20x microscopic objective. Quantified images comprised of 20 different focal layers merged together. Unilateral cell counts of TH and CTb tracer expression were analyzed in VTA spanning four levels across the anterior-posterior plane (AP: −4.92, −5.04, −5.28, −5.40) by one observer.

### Drugs

Methamphetamine HCl (#M8750, Sigma-Aldrich) was dissolved in 0.9% saline (Hospira, Lake Forest, IL, USA) and self-administered intravenously at a dose of 0.1 mg/kg/infusion. Lithium chloride (LiCl; Sigma-Aldrich, IL) was dissolved in water to a concentration of 0.15M and administered intraperitoneally (10 mL/kg).

### Behavioral procedures

Rats were food restricted and maintained to ∼85% of their free-feeding body weight unless otherwise specified. All behavioral testing was conducted in operant chambers from Med-Associates. Each chamber was equipped with a reward magazine that held a receptacle for food pellets and a receptacle for liquid rewards, two retractable levers located to the left and right of the magazine, two stimulus lights (one positioned above each lever), a house light on the opposite chamber wall of the magazine and levers, a pellet dispenser, a syringe pump outside of the chamber, and a swivel attached to a steel tether shielding the drug line. Med-PC V software was used to program and control all hardware. For experiments involving optogenetic techniques, two armored fiber-optic patch cords were connected to a dual-connection rotary joint commutator (Doric Lenses, Quebec, Canada) connected to high-powered DPSS lasers (532nm or 473nm; Shanghai Laser and Optics Century Co., Shanghai, China), which were controlled by Med Associates software. Light leakage from laser output was covered using 5-cm long black shrink-tube shielding over the connected patch cord and cannula ferrules. Chamber contexts differed in flooring, wall-lining, and room location between self-administration procedures, outcome-specific Pavlovian-to-instrumental procedures, and intracranial self-stimulation.

### Magazine Training

Prior to Pavlovian conditioning rats first received one 30-minute session consisting of 30 trials with a variable 60-second intertrial interval (ITI). For each trial, either one 45-mg sucrose pellet or one 0.2 mL bolus of 15% maltodextrin was randomly dispensed into the magazine.

### CS+/CS-Pavlovian Conditioning

Sessions consisted of 12 trials with a variable 6-minute ITI, with one 10s auditory cue (click or white noise) followed by two sucrose pellets (CS+) and the alternative 10s auditory cue (white noise or click) not resulting in pellets (CS-). For VTA_DA_→LH inhibition during conditioning, represented in **Figure 3D**, green laser light was delivered (532 nm, 16-18mW) at the time of pellet delivery, beginning 0.5-sec prior to cue offset and terminating 2-sec after cue offset.

### Blocking Procedure

Rats first received eight sessions of Pavlovian conditioning consisting of 8 trials with a variable 3-minute ITI to acquire two distinct visual cue-pellet associations. Cues (flashing cue lights or steady house light; counterbalanced) were presented for 30 seconds followed by a 1-sec gap before one pellet (45-mg sucrose or 45-mg grain, counterbalanced) was delivered into the magazine. The subsequent four sessions of conditioning introduced novel auditory cues (click or white noise, counterbalanced) each to be presented concurrently with each of the visual cues, followed by the same reward deliveries as before. For one of the cue-pellet pairings, blue light (473nm; 14-16mW;1s; 20Hz; 5ms pulse duration, 45ms interval ^41, 42^) was delivered into the brain at the time of pellet reward, represented in **Figure 4E**. To test whether learning was facilitated by optogenetic stimulation, rats received a probe test consisting of 8 presentations of auditory cues alone without rewards or laser separated by a variable ITI averaging 3 minutes.

### Lithium Chloride-induced Reinforcer Devaluation

Rats were first habituated to the devaluation context by placing them each individually in empty cages in a separate behavioral room from conditioning. Rats received two 30-minute sessions of habituation before being returned to their home cages. Following the last day of habituation, rats were then given 3 daily pairing of the pellets and LiCl, which consisted of 30-minute access to consume 10 grams of pellets immediately followed by intraperitoneal injections of LiCl. Six hours after injection, rats were given their normal home chow to avoid any pairing of their normal diet with LiCl-induced sickness. For non-devalued controls, rats received LiCl injections and were given access to the pellets in their home cages six hours later. Rats were allowed to recover from immediate LiCl effects across 24 hours before being relevant tests. To ensure conditioned aversion to the pellet was present at the time of test, rats received a consumption test conducted immediately after their probe test such that all rats had 10 minutes of free access to pellets in the devaluation cages without subsequent injections.

### Conditioned Reinforcement

Four 30-minute sessions of conditioned reinforcement were conducted in which two levers, never experienced before, were inserted into the behavior chamber. Pressing on one lever (left or right; counterbalanced) produced a 10-sec presentation of CS+ while pressing the other produced the CS-cue. Aligned with the onset and offset of both cues, green laser light was delivered for the entire cue duration (532 nm, 16-18mW). Rats that did not press the lever during these tests were removed from analyses.

### Self-Administration Procedures

Rats received 14 daily self-administration sessions using an adapted procedure ^75, 77, 112^. Animals were trained on increasing fixed-ratio (FR) reinforcement schedules. An FR-1 schedule was used for the first 6 days, followed by an FR-3 schedule for the next 4 days, and then 4 days on an FR-5 schedule ^76^. Sessions consisted of three 1-hour “ON” periods where two levers were inserted into chambers with house light illumination interleaved with two 15-minute “OFF” periods where the levers would retract and the house light was turned off. Across training sessions, rats learned that pressing one lever (“active”) would result in reward and pressing the opposite lever (“inactive”) would have no programmed consequences. Lever designation was counterbalanced across all animals. During the “ON” periods, the Meth group could earn 0.1 mL I.V. infusions of methamphetamine per reward while the Control group could earn two 45-mg grain pellets delivered into the magazine per reward. A 40-second time-out period was initiated in between reward deliveries such that the house light would turn off and both levers would retract before being reinserted back into the chamber and the house light turning back on to signal reward availability. The maximum number of rewards that could be earned per “ON” period for Meth and Control groups was 20 drug infusions or 40 grain pellets, respectively. If this reward limit was reached before the hour was up, the period would complete the remainder of time as an “OFF” period, in which rewards were no longer available to be earned, both levers were retracted, and the house light would turn off, in addition to the subsequent, official 15-minute “OFF” period. Otherwise, should the full hour for the “ON” period elapse before the animal could earn the maximum number of rewards, the session would automatically shift into the 15-minute “OFF” period. Following conclusion of self-administration training, all rats were subject to a 3-week abstinence period where they would remain in their home cages. Rats were taken off food restriction and given *ad libitum* home chow. This abstinence period would allow the Meth group to recover from any somatic symptoms of drug withdrawal that could potentially confound their subsequent learning and performance.

### Outcome-specific Pavlovian-to-Instrumental Transfer

#### Pavlovian Conditioning

All procedures for PIT training and tests were adapted from previous work ^78^. Rats first received eight 60-minute conditioning sessions. During these sessions, two distinct auditory cues (click or white noise) were presented for 2 minutes in a pseudorandom order (4 presentations per cue) with a variable ITI averaging 5 minutes. Each cue was paired with one of two outcomes (sucrose pellets or maltodextrin) that were delivered on a random 30-second schedule across cue presentation. Cue-outcome pairings were counterbalanced across all animals. Entries made into the magazine were used to measure performance and separated by responses during cue presentation (conditioned stimuli; CS) and a 2-minute interval prior to cue onset (baseline).

#### Instrumental Training

Following Pavlovian conditioning, rats received eight training sessions increasing in random-ratio (RR) reinforcement schedules (2x FR1, 3x RR5, 3x RR10) in which they could earn delivery of each of the two outcomes via lever-pressing. One lever was inserted into the chamber per 10-minute trial, alternating levers between trials (4 trials total). Trials were separated by a 2.5m ITI. This timeout period would initiate if: 1) the maximum number of reward outcomes (20 sucrose pellets or 20 boluses of maltodextrin) for the trial was reached or 2) the full 10 minutes for the trial had elapsed. No additional stimuli were present during the session. Lever-outcome pairings were counterbalanced across animals and across their cue-outcome pairings.

#### Transfer Test

Rats were then tested for outcome-specific Pavlovian-to-instrumental transfer (PIT). Both levers were available, but no outcomes delivered. The session began with an 8-minute extinction period to help extinguish lever-press responses produced by the context. Following this, each of the 2-minute auditory cues were presented (4 presentations each) with a fixed ITI of 5 minutes in the following order: click-noise-noise-click-noise-click-click-noise. Entries made into the magazine and lever presses made were recorded, both in the presence of the cue (CS) and in the 2 minutes prior to cue onset (baseline). Lever presses were used to measure PIT performance by separating out responses based on whether the outcome previously associated with a lever corresponded with the same outcome as the cue presented on any given trial (“Same”) or with the opposite outcome (“Diff”).

#### Intracranial self-stimulation

Rats received six 30-minute ICSS sessions defaulted to an FR-1 training schedule unless otherwise stated. Session blocks consist of the average of two individual sessions. Two levers were inserted into the chamber in which pressing on one delivered optogenetic stimulation of the respective terminal projections (“active”) and pressing on the other had no programmed consequences (“inactive”) (counterbalanced). Stimulation consisted of 1- or 2-second trains of blue light [473nm, 14-16mW, 20Hz (LH→VTA: 10ms pulse duration, 40ms interval ^113^; VTA→LH: 5ms pulse duration, 45ms interval), 50Hz (5ms pulse duration, 15ms interval ^41, 42^)].

#### Statistical Analyses

Statistical analyses were conducted using SPSS 28 IBM statistics package. Analyses were conducted using a mixed-design repeated-measures analysis of variance (ANOVA), and t-tests where appropriate. One-tailed tests were used for results with an a priori directional hypothesis at alpha level, *p*=0.1 ^114^. When homogeneity of variances could not be assumed, the more conservative Welch’s t-test was used for analysis. Data for all probe tests and the PIT test were analyzed including all trials of all test sessions. Self-administration and PIT data were correlated using Pearson’s correlations on the change in ratio of active lever presses relative to total number of presses (i.e., active / active + inactive) averaged across the last 3 sessions of self-administration compared to the first 3 sessions, with the ratio of “Same” and “Diff” responses during PIT tests (i.e., Same / Diff). The ICSS data from the self-administration experiment were analyzed across training sessions using log (x+1) transformed data of active and inactive lever presses as the standard deviation was found to increase with the mean ^114, 115^. A formal post-hoc power analysis was conducted on the data elicited from the self-administration x PIT experiment using G*Power 3.0 to estimate the power we had achieved. Here, we used the partial ƞ2 (∼0.62) from our analyses from the PIT test following self-administration to calculate the power (1-β), revealing an estimated power of 0.993 with a type 1 error rate (α) below 0.05. Thus, our sample size for the experimenter-administered methamphetamine control experiment were based on these results. Sample sizes for all other experiments were based on previous work using similar behavioral and neuroscience techniques ^18, 19, 42, 116^.

