## Supplemental Data for "A novel hypothalamic-midbrain circuit for model-based learning"

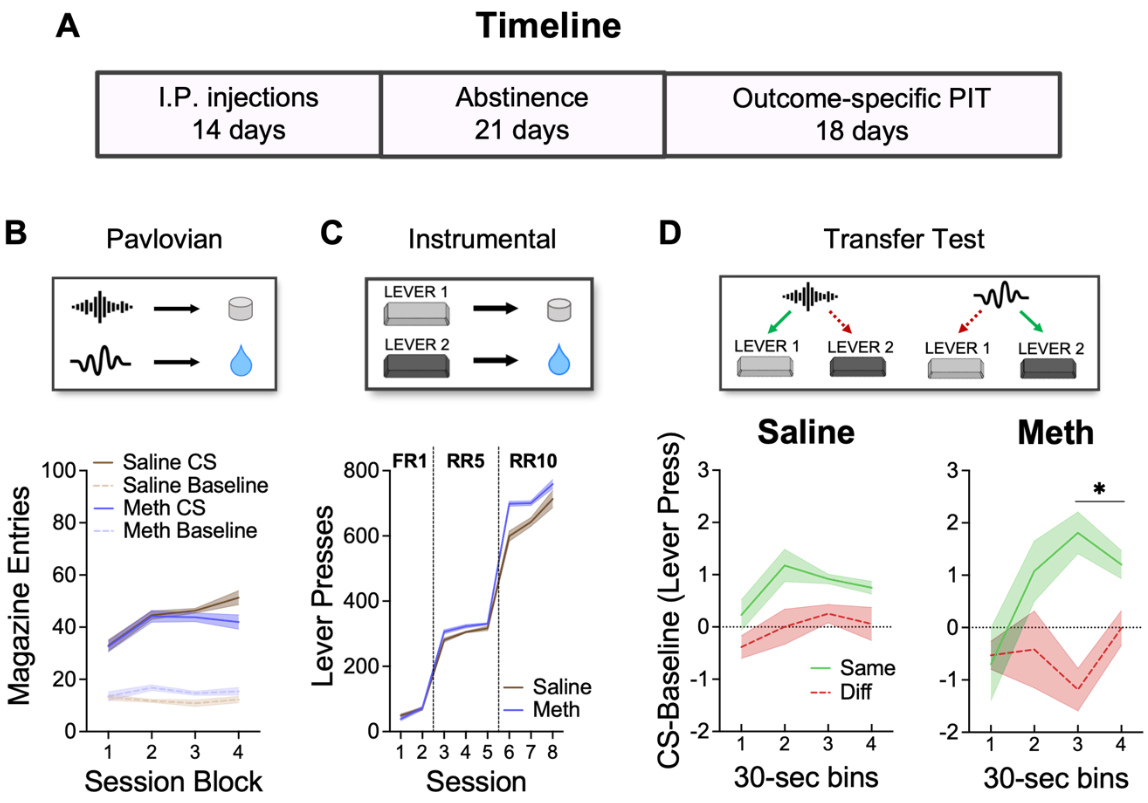


**Supplemental Figure 1. Experimenter-administered methamphetamine also enhances specific PIT.** (A) Experimental timeline. Rats were divided into two treatment groups, with one receiving methamphetamine injections (methamphetamine, *n*=7; 1 mL/kg, I.P.) and the other receiving equivalent volumes of saline (saline, *n*=8; 1 mL/kg, I.P.). (B) Following 3 weeks of abstinence, all rats received Pavlovian conditioning sessions where two auditory cues were paired with two distinct rewards. Both methamphetamine and saline groups increased responding above baseline across time without between-groups differences (CS vs. pre-CS: $F_{(1,13)}$ = 182.768, $p$<0.001; session: $F_{(3,39)}$ = 10.862, $p$<0.001; CS vs. pre-CS x session: $F_{(3,39)}$ = 10.473, $p$<0.001; group: $F_{(1,13)}$ < 0.001, $p$=0.996). (C) Rats were then given instrumental training where two lever presses led to the two distinct rewards. Both groups acquired instrumental responses for the rewards without between-groups differences (session: $F_{(7,91)}$ = 300.031, $p$<0.001; group: $F_{(1,13)}$ = 1.466, $p$=0.247; session x group: $F_{(7,91)}$ = 1.296, $p$=0.261). (D) Finally, rats received the critical PIT test where the auditory cues were presented. Rats in both the saline and methamphetamine groups show greater responding on the same lever relative to the different lever, demonstrating the specific PIT effect (lever: $F_{(1,13)}$ = 14.935, $p$=0.002). However, rats in the methamphetamine group showed an enhanced specific PIT effect, which was most apparent later in the CS (time x lever x group: $F_{(3,39)}$ = 2.497, $p$=0.037; simple main effect of lever, methamphetamine: $F_{(3,39)}$ =

2.994, $p<0.001$; simple main effect of lever, saline: $F_{(3,39)} = 0.661$, $p=0.117$), mirroring our effects seen with self-administration of methamphetamine. *$p \leq 0.05$, mean (± SEM).

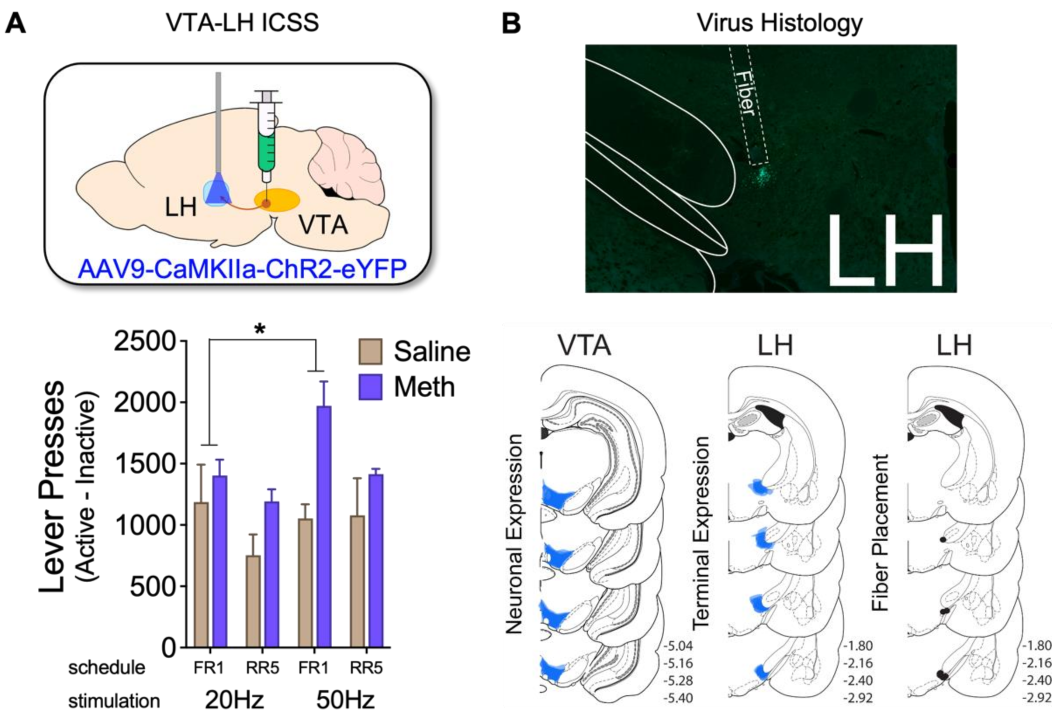


**Supplemental Figure 2. Experimenter-administered methamphetamine sensitizes the VTA → LH pathway.** (A) Rats that received non-contingent methamphetamine (or saline control) injections underwent surgeries to infuse an excitatory channelrhodopsin virus into the VTA with optic fibers implanted over LH. Rats then received sessions of ICSS under low (20 Hz) and high (50 Hz) frequency optogenetic stimulation across increasing schedules of reinforcement. Compared to saline controls, rats in the methamphetamine group show greater self-stimulation, represented as the difference in presses between active and inactive levers (i.e., active – inactive), for VTA terminals in LH when the stimulation condition increased to a higher frequency (lever x stimulation x schedule x group: $F_{(1,6)}$ = 4.226, $p$=0.086; simple main effect of stimulation on FR1, methamphetamine: $F_{(1,6)}$ = 630.000, $p$=0.046; simple main effect of stimulation on FR1, saline: $F_{(1,6)}$ = 138.667, $p$=0.601), suggesting that methamphetamine produced a hypersensitivity to VTA→LH stimulation when stimulation of this pathway shifted to "rewarding" frequencies [79-81]. (B) Unilateral example of bilateral virus expression in the axonal terminals in LH (*top*), and schematics of virus expression in VTA and LH with fiber placements in LH (*bottom*). *$p \leq 0.05$, mean (± SEM).
